# Discovery of Chlorofluoroacetamide-Based Covalent Inhibitors for SARS-CoV-2 3CL Protease

**DOI:** 10.1101/2022.06.05.494897

**Authors:** Yuya Hirose, Naoya Shindo, Makiko Mori, Satsuki Onitsuka, Hikaru Isogai, Rui Hamada, Tadanari Hiramoto, Jinta Ochi, Daisuke Takahashi, Tadashi Ueda, Jose M.M. Caaveiro, Yuya Yoshida, Shigehiro Ohdo, Naoya Matsunaga, Shinsuke Toba, Michihito Sasaki, Yasuko Orba, Hirofumi Sawa, Akihiko Sato, Eiji Kawanishi, Akio Ojida

## Abstract

The pandemic of coronavirus disease 2019 (COVID-19) has urgently necessitated the development of antiviral agents against severe acute respiratory syndrome coronavirus 2 (SARS-CoV-2). The 3C-like protease (3CL^pro^) is a promising target for COVID-19 treatment. Here, we report the new class of covalent inhibitors for 3CL^pro^ possessing chlorofluoroacetamide (CFA) as a cysteine reactive warhead. Based on the aza-peptide scaffold, we synthesized the series of CFA derivatives in enantiopure form and evaluated their biochemical efficiencies. The data revealed that **8a** (**YH-6**) with *R* configuration at the CFA unit strongly blocks the SARS-CoV-2 replication in the infected cells and this potency is comparable to that of nirmatrelvir. The X-ray structural analysis shows that **8a** (**YH-6**) forms a covalent bond with Cys145 at the catalytic center of 3CL^pro^. The strong antiviral activity and sufficient pharmacokinetics property of **8a** (**YH-6**) suggest its potential as a lead compound for treatment of COVID-19.

## Introduction

The outbreak of the novel coronavirus disease 2019 (COVID-19), caused by severe acute respiratory syndrome coronavirus 2 (SARS-CoV-2),^1,2^ poses a severe threat to global public health and economy. Although efficacious vaccines are now being administered worldwide, the development of antiviral agents against SARS-CoV-2 is urgently needed to reduce the cases and also severity of symptoms and fatality rates.^3,4^ SARS-CoV-2 has a single-stranded genomic RNA that encodes polyproteins pp1a and pp1ab. Two cysteine proteases, 3C-like protease (3CL^pro^, also known as main protease, M^pro^) and papain-like protease (PL^pro^), are excised from the polyproteins and further digest the polyproteins into nonstructural proteins, including crucial components of the viral replication–translation machinery. Although both proteases are essential for viral replication, the predominant role of 3CL^pro^ in polyprotein processing and the lack of a human homolog has led to extensive studies to develop an attractive drug target against SARS-CoV-2.^5,6^

3CL^pro^ recognizes glutamine and a hydrophobic amino acid residue at the S1 and S2 pockets in the active site, respectively, and cleaves amide bonds primarily within the Leu-Gln↓ (Ser, Ala, Gly) sequence (↓ : cleavage site).^7^ This substrate specificity is conserved across other coronaviruses, including SARS-CoV-1 and middle east respiratory syndrome coronavirus (MERS-CoV).^8,9^ Therefore, initial efforts to develop SARS-CoV-2 3CL^pro^ inhibitors are extensively guided by the previous molecular designs, especially those targeting SARS-CoV-1 3CL^pro^.^4,5^ Most of them are peptidomimetics based on the Leu-Gln sequence, combined with an electrophilic warhead, such as α-ketoamide,^10^ aldehyde,^11–14^ ketone,^15–18^ vinyl sulfone,^7^ and nitrile,^19-21^ for the covalent capture of the sulfhydryl group of the catalytic Cys145. Nirmatrelvir (PF-07321332), developed by Pfizer, is one of the most advanced compounds in this category and has been recently approved as an oral drug in combination with ritonavir (Fig. 1A).^20^ Other than peptidomimetics, several Ugi multicomponent reaction (MCR)-generated dipeptidic compounds have also been reported as the inhibitors against 3CL^pro^ (Fig. 1B).^22–26^ Very recently, Shionogi has reported S-217622 as non-peptidic and non-covalent inhibitor for 3CL^pro^.^27^

**Figure 1.**
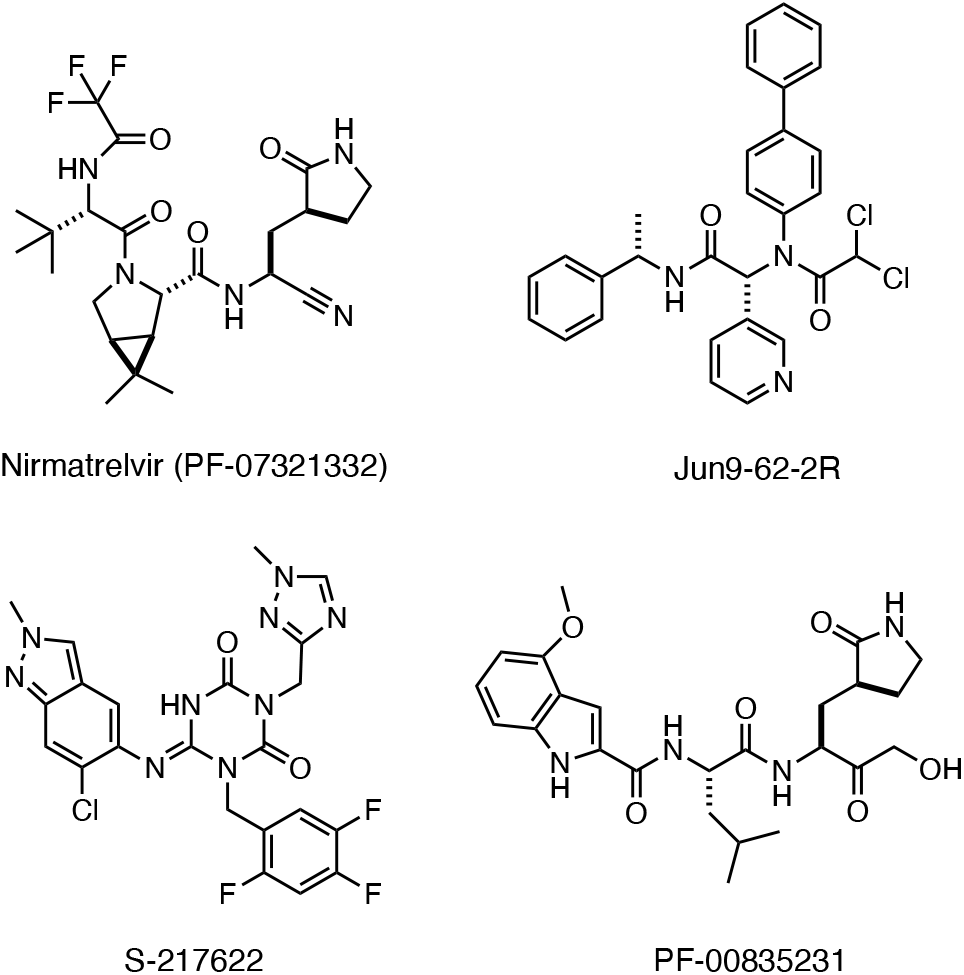
Structures of the SARS-CoV-2 3CL^pro^ inhibitors.

Covalent inhibition of proteins has been a powerful strategy to achieve potent and sustained pharmacological efficacy of small molecule inhibitors.^28^ We have recently introduced α-chlorofluoroacetamide (CFA) as a weakly reactive, cysteine-directed warhead for covalent targeting of proteins.^26,29–31^ We found that the CFA derivatives exhibited higher selectivity in covalent modification of tyrosine kinases than the structurally related acrylamide-based inhibitors. We also demonstrated that the CFA–thiol adduct is hydrolyzed under neutral aqueous conditions to reversibly generate an unmodified cysteine, which could contribute to high target selectivity by eliminating the solvent-exposed off-target labeling. We envisioned that these desirable features of CFA also would be useful in development of selective covalent inhibitor for 3CL^pro^.

In the paper, we report the development of irreversible inhibitor for 3CL^pro^ of SARS-CoV-2 using CFA as a cysteine-reactive warhead. Introduction of a CFA unit into aza-peptide scaffold provided the covalent inhibitors that show potent inhibitory activity against 3CL^pro^. Synthesis of the enantiopure CFA derivative and screening of their biological activity led to identify **YH-6** with a (*R*)-CFA unit as a promising lead compound for treatment of COVID-19.

## Results and discussion

### Synthesis of Inhibitors

We designed the CFA-based covalent inhibitors based on the structure of PF-00835231, which has been reported by Pfizer as a ketone-based covalent inhibitor for 3CL^pro^. The X-ray structural analysis revealed that PF-00835231 generates a tetrahedral carbinol complex with the catalytic Cys145 of 3CL^pro^ at its carbonyl center. We envisioned that replacement of the α-hydroxyl ketone group of PF-00835231 to CFA unit would provide a new class of reversible covalent inhibitor against 3CL^pro^ with high target selectivity. For this purpose, we employed the azapeptide scaffold,^32^ which realizes introduction of a CFA unit with minimum structural perturbation.

To construct the chiral architecture of the CFA-based inhibitors, we streamlined the synthetic route, which utilizes the chiral building blocks and chromatographic chiral separation. The synthetic route of the inhibitors are shown in Scheme 1 and 2. Carboxybenzoyl (Cbz) protection of the commercially available 2-pyrrolidone gave **1** according to the reported method.^33,34^ Formylation of **1** using *tert*-Butoxybis(dimethylamino)methane (Bredereck’s reagent)^35,36^ followed by the treatment with aqueous hydrochloric acid yielded **2** in quantitative yield. Conjugation of **2** and *N*-protected L-leucine hydrazides **3** was conducted by reductive amination using dimethylamine borane and p-toluenesulfonic acid to yield **4** in 68% yield. Protection of the amino group of **2** by *tert*-butoxycarbonyl (Boc) group to give in 79% yield, and the subsequent deprotection of the Cbz group by palladium-catalyzed hydrogenolysis gave **6** in 95% yield as a diastereomixture. For the separation of the diasteromers, we employed medium pressure liquid chiral column chromatography (chiral MPLC). The optimized separation conditions using a CHIRALFLASH^®^ IC column (Daicel) enabled us to obtain a multi-hundred milligram of enantiopure **(*S***,***S*)-6** and **(*S***,***R*)-6**.

**Scheme 1.**
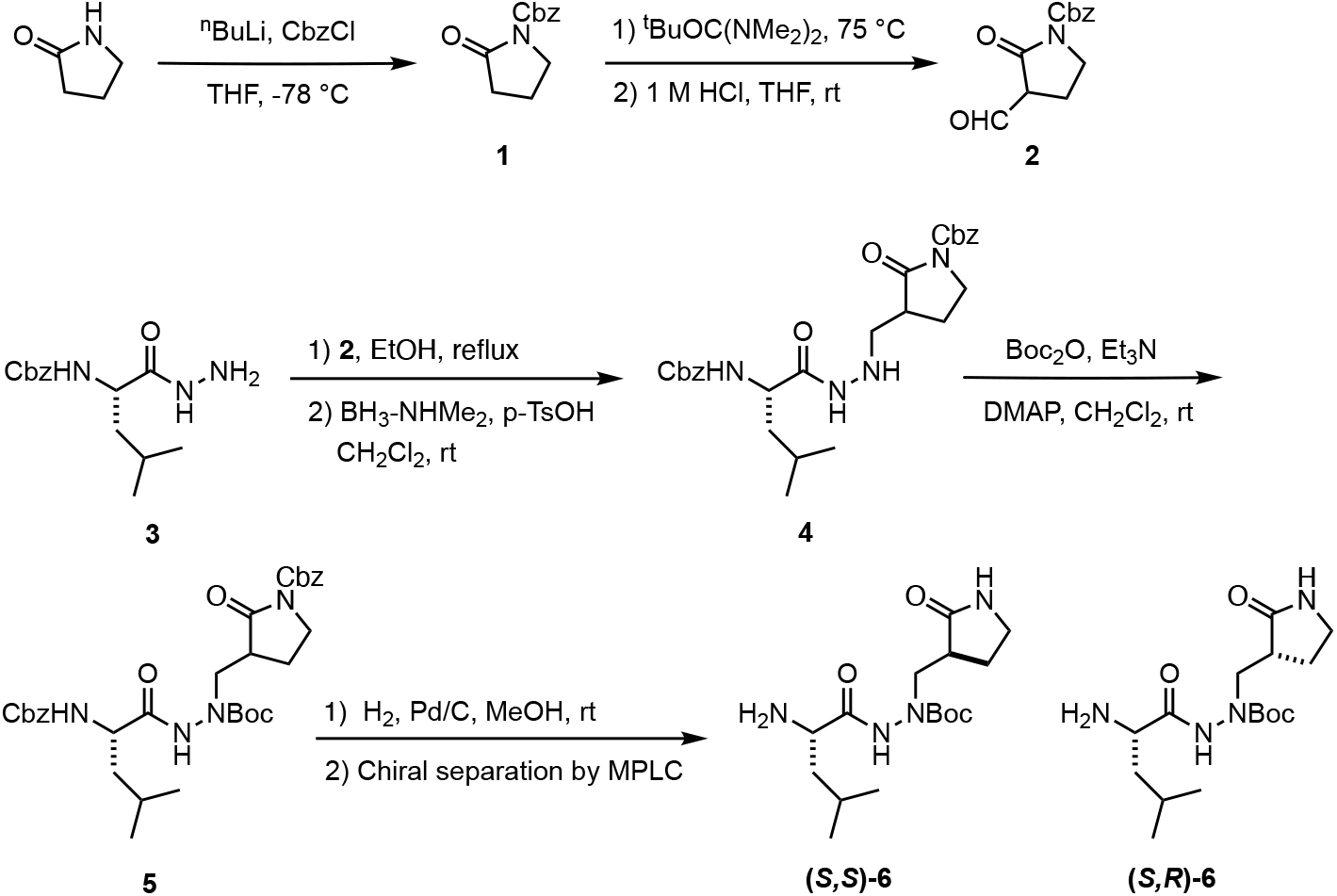
Synthesis of chiral intermediate **6**.

The enantiopure intermediate **(*S***,***S*)-6** was conjugated with the indole 2-carboxylate (R = F or OMe) using COMU as a coupling reagent to give **7a** or **7b** (Scheme 2). After deprotection of the Boc group, **7a** (R = F) was converted to **8a** (**YH-6)** or **9** by the conjugation reaction with (*R*)- or (*S*)-chlorofluoroacetic acid^26^, respectively. The conjugate reaction of **7b** (R = OMe) with (*R*)-chlorofluoroacetic acid also provided **8b**. The same synthetic procedure was applied to the enantiopure intermediate **(*S***,***R*)-6** to obtain CFA derivatives **10a-b** and **11**. The absolute chiral configurations of these compounds were determined based on the X-ray structural analysis of **8b**, which was assigned to be a (*R*)-CFA and (*S*)-pyrrolidone stereoisomer (Figure S1, Table S1).

**Scheme 2.**
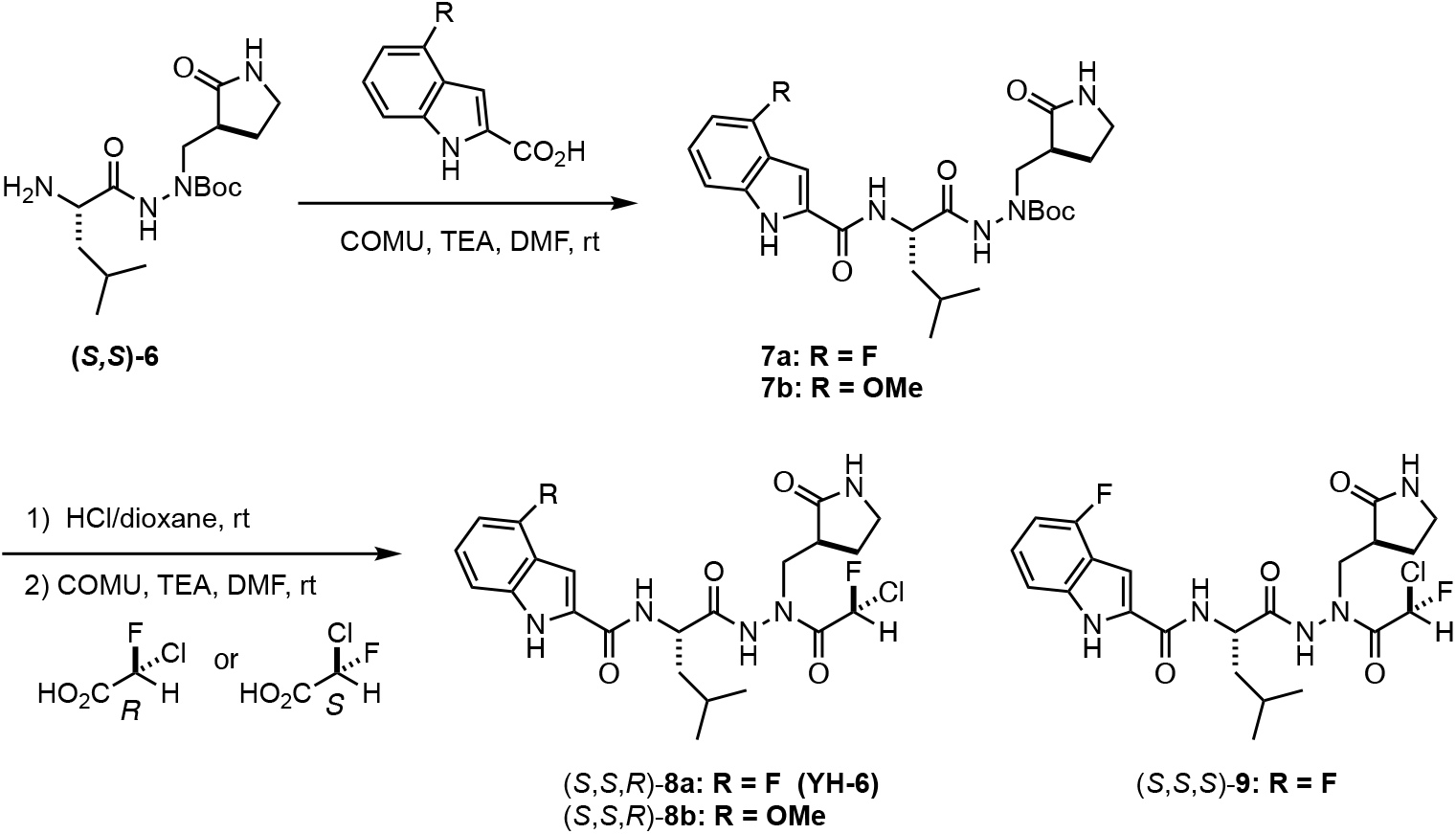
Synthesis of the CFA-based inhibitors (**8a-b, 9**).

### Biological Evaluation

The inhibitory activity of the CFA derivatives was first assessed by the in vitro enzymatic assay using the recombinant SARS-CoV-2 3CL^pro^ and the fluorogenic peptide substrate Ac-Abu-Tle-Leu-Gln-(4-methylcoumaryl-7-amide).^7^ We found that **8a** (**YH-6**) exhibits the strongest inhibitory activity among the tested CFA derivatives. The IC_50_ value of **8a** (**YH-6**) was 3.8 nM, which is slightly larger than that of nirmatrelvir (IC_50_ = 0.8 nM) but comparable to that of PF-00835231 (IC_50_ = 3.7 nM). Interestingly, the activity of **8a** (**YH-6**) was stronger than the corresponding (*S*)-CFA isomer **9** (IC_50_ = 21 nM). Further, the acetyl analogue **13**, which lacks a CFA unit, showed a very weak (IC_50_ = 354 nM). These results suggest that the (*R*)-configuration of the CFA unit of **8a** (**YH-6**) is crucial for the strong inhibition toward 3CL^pro^ via the efficient covalent bond formation with Cys145 of 3CL^pro^. Regarding the chirality of the pyrrolidone unit, (*S*)-2-pyrrolidone **8a** (**YH-6**) exhibited a slightly higher inhibitory activity than the corresponding (*R*)-2-pyrrolidone **10a** (IC_50_ = 7.8 nM). The same stereoselective inhibition was also observed in the case of the 4-methoxyindole derivatives, in which (*S*)-2-pyrrolidone **8b** (IC_50_ = 6.3 nM) is more potent than (*R*)-2-pyrrolidone **10b** (IC_50_ = 15.6 nM). These data suggest that the pair of (*R*)-CFA and (*S*)-2-pyrrolidone configurations of **8a** (**YH-6**) is a matched chirality preferred for the potent inhibition toward 3CL^pro^. Indeed, **11** with the opposite (*S*)-CFA and (*R*)-2-pyrrolidone configurations showed a week inhibitory activity (IC_50_ = 178 nM). We confirmed that **8a** (**YH-6**) shows the negligible inhibition activities against human cysteine proteases such as cathepsin L (IC_50_ > 10 µM) and B (IC_50_ > 10 µM) in the in vitro enzymatic assays.

The antiviral activity of the inhibitors was evaluated based on the inhibition of the cytopathic effects of SARS-CoV-2 elicited in the infected cells (Table 2). We found that **8a** (**YH-6**) and **8b** exhibited the strong antiviral activities in 293T-ACE2-TMPRSS2 (293TAT) cells infected with various SARS-CoV-2 strains including wild type (WT), alpha, beta, omicron BA.1 and omicron BA.2. This result is quite reasonable because no significant mutation have been reported near the catalytic site of 3CL^pro^ in the strains reported to date. The EC_50_ values of **8a** (**YH-6**) and **8b** reached to a single digit nanomolar range, which is comparable to those of nirmatrelvir and much lower than those of PF-00835231 and N^4^-Hydroxycytidine (NHC, active form of molnupiravir as an RNA dependent RNA polymerase inhibitor). The activities of **8a** (**YH-6**) and **8b** were stronger than those of the corresponding diastereomers **10a** and **10b**. This result is consistent with that of the in vitro enzymatic assay using the recombinant 3CL^pro^ (Table 1). It was found that that the antiviral activities of the CFA-based inhibitors largely decreased in VeroE6T cells infected with wild type SARS-CoV-2. However, the potencies were effectively recovered by the presence of CP-100356 (0.5 µM) as an inhibitor of p-glycoprotein (p-gp).^37,38^ These pronounced potency shifts suggest that the CFA-based inhibitors act as the substrates of the p-gp efflux pump, as similar to PF-00835231.^15^ Although this would be a potential concern in drug development, the recent paper reported that such p-gp mediated efflux does not negatively impact the PF-00835231 efficacy in human polarized airway epithelial cultures.^39^ Further in vivo study on the antiviral activity of **8a** (**YH-6**) need to be pursued. The antiviral activities of **8a** (**YH-6**) and **8b** were further tested in the 293TAT and 293T-DPP4 cells infected with other viruses such as SARS-CoV, MERS-CoV, and HCoV-OC43. Interestingly, **8a** (**YH-6**) and **8b** displayed the stronger activities (EC_50_ = 1.4 ∼ 3.6 nM) against MERS-CoV, and HCoV-OC43 as compared to SARS-CoV-1 and the SARS-CoV-2 variants, suggesting a broad utility of the CFA-based 3CL^pro^ inhibitors against the various corona virus infections.

**Table 1.** Enzymatic inhibitory activity against SARS-CoV-2 3CL^pro^.

| | 3CL<sup>pro</sup> $IC_{50}$<br>(nM)<sup>a</sup> |
| --- | --- |
| **8a** | 3.8 |
| **8b** | 6.3 |
| **9** | 21.0 |
| **10a** | 7.8 |
| **10b** | 15.6 |
| **11** | 355 |
| **12** | 178 |
| PF-00835231 | 1.8 |
| nirmatrelvir | 0.81 |

<sup>a</sup> $IC_{50}$ value was obtained by the plot of the average of triplicate experiments.

**Table 2.** Antiviral activity against various SARS-CoV-2 strains in cytopathic effect inhibition assay with 293TAT cell.

| | 293TAT $EC_{50}$ (nM)<sup>a</sup> | | | | |
| --- | --- | --- | --- | --- | --- |
| | WT | $\alpha$ | $\delta$ | Om BA.1 | Om BA.2 |
| **8a** | 21.2 ± 2.7 | 13.8 ± 1.3 | 7.57 ± 2.59 | 9.01 ± 2.45 | 17.1 ± 2.5 |
| **8b** | 24.4 ± 1.3 | 16.2 ± 2.5 | 8.79 ± 1.51 | 10.1 ± 2.3 | 18.3 ± 3.0 |
| **10a** | 141 ± 15 | 92.3 ± 3.4 | 44.5 ± 17.5 | 58.7 ± 5.9 | 131 ± 24 |
| **10b** | 119 ± 8 | 103 ± 1 | 50.0 ± 22 | 58.8 ± 4.3 | 121 ± 29 |
| PF-00835231 | 915 ± 53 | 1604 ± 44 | 692 ± 325 | 775 ± 254 | 832 ± 52 |
| nirmatrelvir | 21.7 ± 3.6 | 56.9 ± 7 | 30.2 ± 19.8 | 21.1 ± 3.1 | 21.7 ± 2.8 |
| NHC<sup>c</sup> | 446 ± 36 | 1222 ± 201 | 351 ± 38 | 574 ± 273 | 336 ± 168 |

<sup>a</sup> Data represent mean ± standard deviation of repeated experiments (n = 3 or 4). <sup>c</sup> active form of molnupiravir (RNA dependent RNA polymerase inhibitor).

**Table 3.** Antiviral activity against various SARS-CoV-2 strains in cytopathic effect inhibition assay with VeroE6/TMPRSS2 cells.

| | VeroE6T $EC_{50}$ (nM)<sup>a</sup> | |
| --- | --- | --- |
|  | WT | WT+CP<sup>b</sup> |
| **8a** | 6551 ± 1606 | 59 ± 4.8 |
| **8b** | 20840 ± 5352 | 69.7 ± 0.9 |
| **10a** | >50000 | 423±70 |
| **10b** | >50000 | 389±67 |
| PF-00835231 | >50000 | 509±37 |
| nirmatrelvir | 5205 ± 789 | 27.3 ± 2.3 |
| NHC<sup>c</sup> | 204 ± 69 | 238 ± 17 |

<sup>a</sup> Data represent mean ± standard deviation of repeated experiments (n = 3 or 4). <sup>b</sup> CP-100356.
<sup>c</sup> active form of molnupiravir (RNA dependent RNA polymerase inhibitor).

**Table 4.** Antiviral activity against various corona viruses in cytopathic effect inhibition assay with 293TAT and 293T-DPP4 cell.

|  | EC50 (nM)<sup>a</sup> |  |  |
| --- | --- | --- | --- |
|  | 293TAT | 293T-DPP4 | 293TAT |
|  | SARS-CoV | MERS-CoV | HCoV-OC43 |
| **8a** | 59.3 ± 14.5 | 4.72 ± 0.22 | 1.67 ± 0.39 |
| **8b** | 64.4 ± 15.3 | 5.01 ± 0.69 | 3.18 ± 0.18 |
| PF-00835231 | 3236 ± 148 | 700 ± 35 | 929 ± 71 |
| nirmatrelvir | 70.2 ± 16.2 | 24.3 ± 1.4 | 39.8 ± 7.0 |
| NHC<sup>b</sup> | 1556 ± 189 | 1318 ± 664 | 1194 ± 134 |

<sup>a</sup> Data represent mean ± standard deviation of repeated experiments (n = 3 or 4). <sup>b</sup> active form of molnupiravir (RNA dependent RNA polymerase inhibitor).

We conducted a preliminary pharmacokinetics study in mice to estimate ADME property of the CFA derivatives. When **8a** (**YH-6**) was orally administered to mice (100 mg/kg), its plasma concentration reached to 15.2 µM after 1 h and decreased to 0.40 µM after 6 hr. On the other hand, oral administration test of **8b** was failed to due to its toxic effect on mice at the same dose. In intravenous administration of **8a** (**YH-6**) (10 mg/kg), its plasma concentration reached to 9.3 µM after 1 hr and decreased to 0.33 µM after 6 hr. Based on these data, the bioavailability (BA) of **8a** (**YH-6**) was roughly estimated to be 11%. We also confirmed that concentration of **8a** (**YH-6**) in mouse lung tissue reached to 78 and 63 nmol/mg after 6 hr of the oral (100 mg/kg) and the intravenous (10 mg/kg) administration, respectively (Figure S2). In the cell viability assays using the SARS-CoV-2 infected 293TAT cells, EC_50_ values of **8a** (**YH-6**) were slightly lower in the presence of 2% human serum albumin (HSA) (Table S2), implying the low binding property of **8a** (**YH-6**) to HSA in plasma. Overall, these inhibition activity and PK data suggest that **8a** (**YH-6**) has strong potential for further drug development.

### X-ray Crystal Structure

To clarify the molecular basis of the inhibition of 3CL^pro^ by the CFA derivatives, we determined the X-ray structure of 3CL^pro^ in complex with **8a** (**YH-6**) at 1.6 Å resolution (PDB code: 7XAR, Table S3). The crystallographic asymmetric unit comprises the 3CL^pro^ dimer. Electron density revealed that **8a** (**YH-6**) has an extended conformation throughout the substrate binding channel, in which the side chains of **8a** (**YH-6**) occupy the S1, S2, and S3 binding pockets. In the complex, the CFA unit of **8a** (**YH-6**) forms a covalent bond with the catalytically active Cys145 residue (Figure 4a). The *S*-configuration of the CFA carbonyl center in the complex clearly suggest that the nucleophilic attack of the sulfur atom of Cys145 proceeds through a *S*_N_2 reaction mechanism. The 4-fluroindole group of **8a** (**YH-6**) is covered by Gln189, Thr190, and Ala191 in the neighborhood of the S3 position, which could contribute to the enhanced inhibition activity of **8a** (**YH-6**) by non-polar interactions (Figure 4b). In the S1’ catalytic pocket, the carboxamide oxygen of CFA unit engaged in two hydrogen bonds with the NH group of the main chain of residues Glys143 and Cys145, which could be important to stabilize the reversible covalent bond with the Cys145 residue.

**Figure 3.**
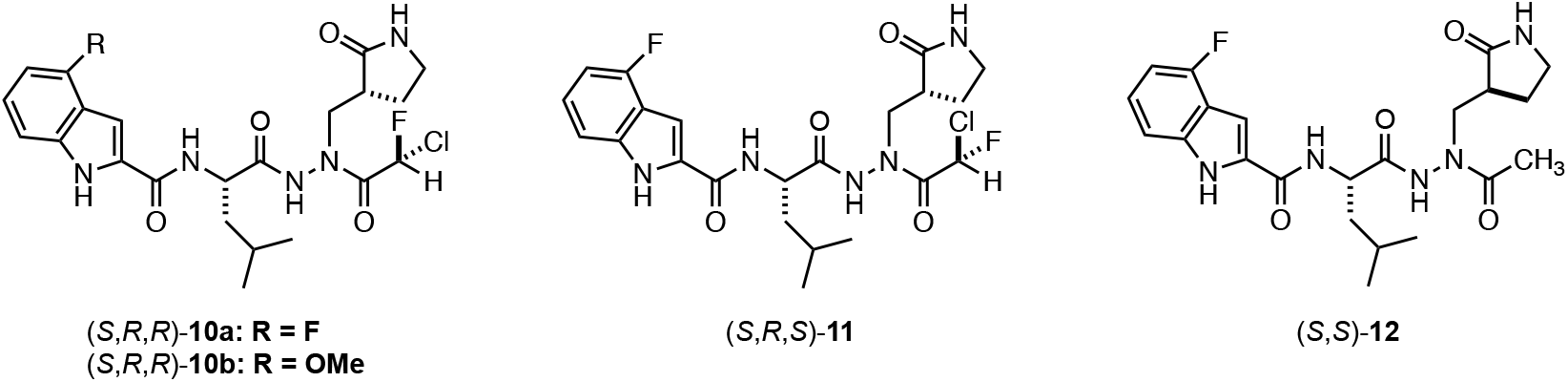
Structures of CFA-based inhibitors (**10a-b, 11**) and acetyl analogue **12**.

**Figure 3.**
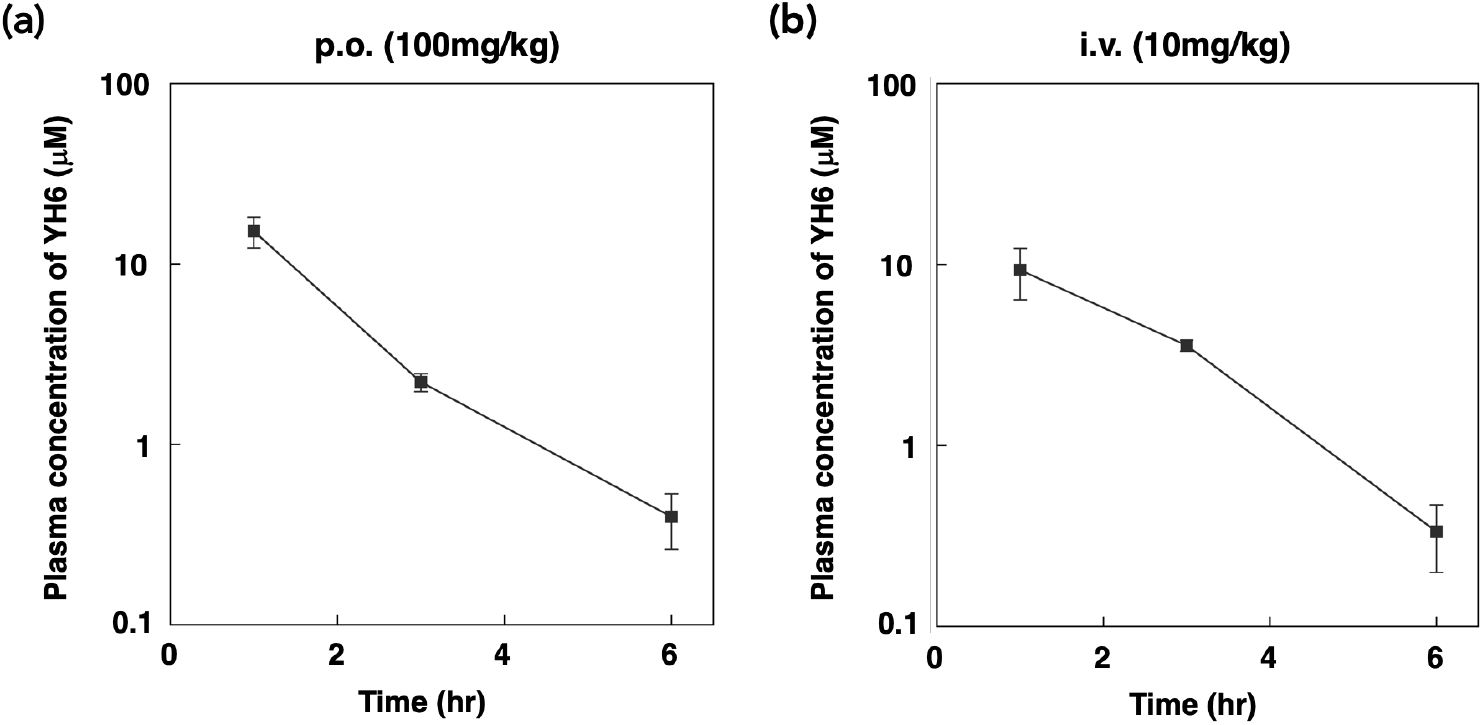
Mean plasma concentration (µM)-time profile of **8a** (**YH-6**) in mice (fasted state) after (a) p.o. (100 mg/kg) and (b) i.v. (10 mg/kg) administration. Data are shown as the mean ± SE (n = 3).

**Figure 4.**
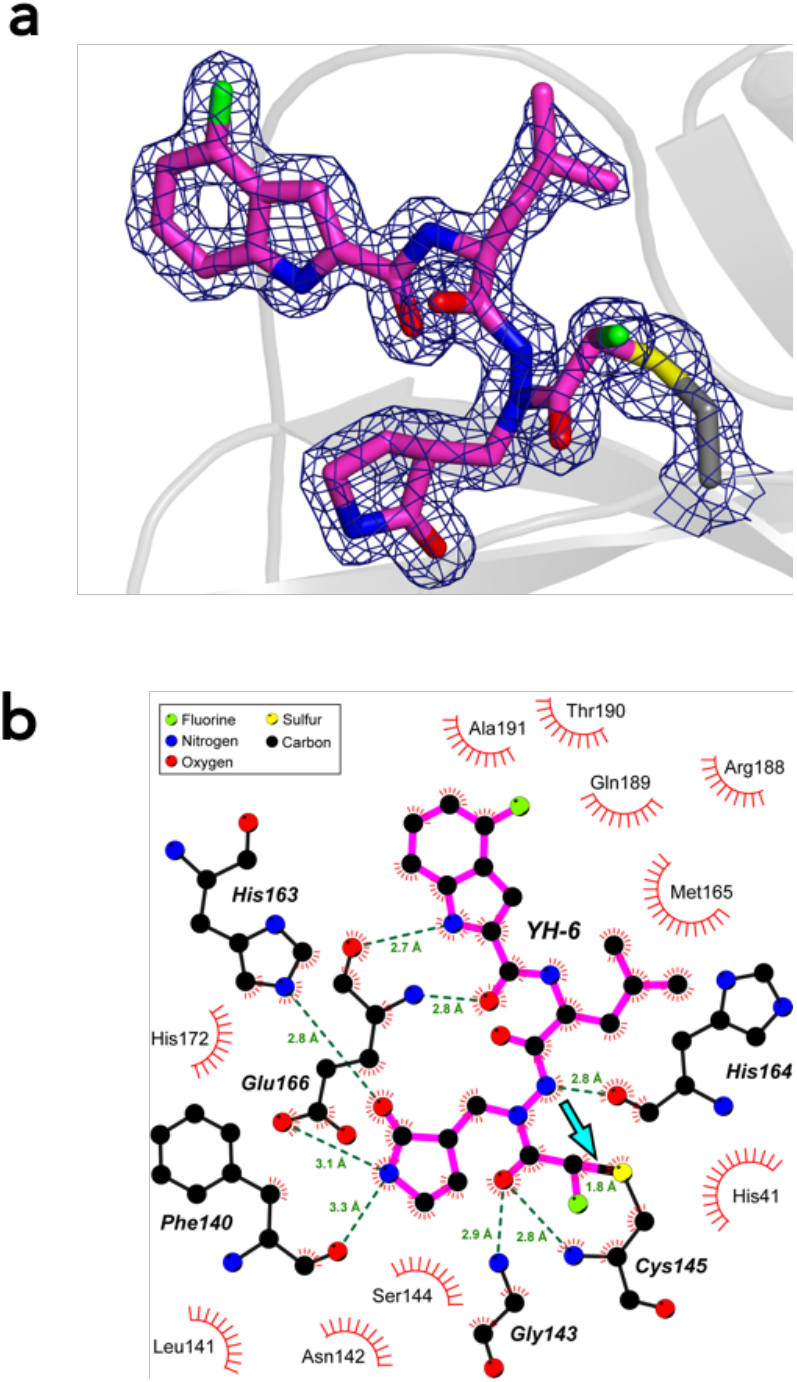
X-ray crystal structure of SARS-CoV-2 M^pro^ in complex with **8a** (**YH-6**). (a) Sigma-A weighted 2*F*_o_ − *F*_c_ electron density map for **8a** (**YH-6**) and Cys145 contoured at 1.5 σ (blue mesh). The refined model of **8a** (**YH-6**) covalently bound to Cys145 of 3CL^Pro^ is shown as magenta, blue, red and green sticks, corresponding to carbon, nitrogen, oxygen and fluorine atoms, respectively. Carbon and sulfur atoms of Cys145 are shown as gray and yellow sticks, respectively. (b) Interaction network between 3C-like protease and covalent inhibitor **YH-6**. The inhibitor is represented by color spheres and magenta lines. Protein residues engaging in H-bonds with **8a** (**YH-6**) are represented by color spheres and black lines. H-bonds are indicated by the green dotted lines. Protein residues engaging in non-bonding interactions with **8a** (**YH-6**) are shown as red spoked arcs. The blue arrow points at the covalent bond between residue Cys145 and the inhibitor, which is shown as a thick solid line, half magenta, half black. The plot was prepared with LigPlus+.^40^

## Conclusion

In summary, we have developed a new class of covalent inhibitor of SARS-CoV-2 3CL^pro^ using CFA as a reactive warhead. The introduction of aza-peptide scaffold and the establishment of the chiral synthetic method allowed us to identify **8a** (**YH-6**) as a potent inhibitor against 3CL^pro^. The findings in the biological assays prompted further preclinical investigation of **8a** (**YH-6**) as a potential agent for treatment of COVID-19. We also believe that the prominent role of CFA chirality in the covalent inhibition of 3CL^pro^ would provide a useful knowledge to design covalent inhibitor with different molecular architecture targeting other disease-associated proteins. Our research is ongoing along this line and will be reported in due course.

## Acknowledgements

We thank Dr. Taisuke Matsumoto (Institute for Materials Chemistry and Engineering, Kyushu University) for the X-ray diffraction experiment. This work was supported by AMED under Grant Number JP20fk0108519 and Platform Project for Supporting Drug Discovery and Life Science Research (Basis for Supporting Innovative Drug Discovery and Life Science Research (BINDS) under Grant Number JP18am0101091. A.O. acknowledges Grant-in-Aid for Scientific Research on Innovative Areas “Chemistry for Multimolecular Crowding Biosystems” (JSPS KAKENHI Grant No. JP17H06349) for its financial support. N.S. acknowledges Grant-in-Aid for Scientific Research B (JSPS KAKENHI Grant No. 19H02854), AMED under Grant Number JP21ak0101121, and Grant for Basic Science Research Projects from the Sumitomo Foundation, for their financial supports. We thank Dr. Koichi Morita (Nagasaki University, Japan) for providing SARS-CoV Hanoi strain, Dr. Bart Haagmans (Erasmus University Medical Center, Netherlands) for providing MERS-CoV EMC2012 strain, Drs. Masayuki Saijyo, Masayuki Shimojima, Mutsuyo Ito and Ken Maeda (National Institute of Infectious Diseases (NIID), Japan) for providing SARS-CoV-2 variants.

## Experimental section

### Fluorescence 3CL^pro^ inhibitory assay

SARS-CoV-2 main protease (3CL^pro^) (100823), human cathepsin B (80001), and human cathepsin L (80005) were purchased from BPS Bioscience (San Diego, CA). The assay was performed in 96-well micro plate using the reaction buffer containing 20 mM Tris-HCl (pH 7.3), 100 mM NaCl, 1 mM EDTA, and 1 mM DTT. A solution of 25 nM SARS-CoV-2 3CL^pro^ incubated with the test compound at the different concentrations (20 µL) for 30 min at ambient temperature. The enzymatic reaction was initiated by addition of the fluorogenic substrate (Ac-Abu-Tle-Leu-Gln-MCA, Peptide Institute 3250-v) (50 µM, 10 µL) and allowed to proceed for 6 h at ambient temperature. The fluorescence intensity of each well was measured using EnSpire multimode plate reader (PerkinElmer) at excitation wavelength of 355 nm and emission wavelength of 460 nm. Percent activity of the inhibition was calculated based on the fluorescence intensity of the control well without inhibitor (100% activity) and that of the blank well without 3CL^pro^ (0% activity). IC_50_ value was calculated by curve fitting analysis using KaleidaGraph (Synergy Software). Triplicate experiments were performed for each data point. Cathepsin enzymatic assays were performed in the reaction buffer containing 20 mM sodium acetate (pH 5.5), 1 mM EDTA, and 5 mM DTT. 750 pM cathepsin B or 750 pM cathepsin L (20 µL) and the compound (125 µM, 20 µL) were added to each well and incubated for 30 min at ambient temperature. The enzymatic reaction was initiated by adding the fluorogenic substrate (Z-Leu-Arg-MCA, Peptide Institute 3210-v) (25 µM, 10 µL) and allowed to proceed for 1 hr at ambient temperature. Percent inhibition was calculated as described above.

### Cells and viruses

Vero-TMPRSS2 cells [Vero E6 cells (ATCC, CRL-1586) stably expressing human TMPRSS2]^41^ were maintained in Dulbecco’s Modified Eagle’s Medium (DMEM) containing 10% FBS. 293T (RIKEN BRC, RCB2202) cells were maintained in high glucose DMEM containing 10% FBS. 293TAT and 293T-DPP4 cells were established by transduction of human TMPRSS2, ACE2 or DPP4 expression cassettes using lentiviral vectors as previously described ^42^ and maintained in high glucose DMEM containing 10% FBS.

SARS-CoV-2 VOC Alpha (strain QK002, lineage B.1.1.7, GISAID: EPI_ISL_768526), Delta (strain TY11-927, lineage B.1.617.2, GISAID: EPI_ISL_2158617), Omicron (strain TY38-873, lineage BA.1, GISAID: EPI_ISL_7418017, BA.2 GISAID: EPI_ISL_9595859) were provided from National Institute of Infectious Diseases, Japan. The working viral stocks were prepared by a passage on Vero-TMPRSS2 cells. SARS-CoV strain Hanoi was kindly provided by Nagasaki University and was amplified in VeroE6 cells. MERS-CoV strain EMC2012 was kindly provided by Erasmus University Medical Center and amplified on VeroE6-TMPRSS2 cells. Human coronavirus strain OC43 (HCoV-OC43) were purchased from ATCC (VR-1558) and was amplified in 293T cells. The titers of the prepared virus stocks were determined as plaque forming unit per ml (pfu/ml) by a plaque assay.

### Cytopathic effect-based cell viability assays

Nirmatrelvir and PF-00835231 were purchased from Selleckchem. Molnupiravir (MPV, also known as EIDD-2801) is a ribonucleoside prodrug of N-hydroxycytidine (NHC) and targets viral RNA polymerase of SARS-CoV-2. NHC was obtained from Angene. The compounds were serially diluted 2-fold increments by culture medium containing 2% FBS and plated on 96-well microplates. The diluted compounds in the plates were mixed with SARS-CoV-2 and cell suspension. Cells in the plates were cultured for 2-3 days, and then exposed with MTT (3-[4,5-dimethyl-2-thiazolyl]-2,5-diphenyl-2H-tetrazolium bromide) (Nacalai Tesque). Cell viability was determined by measurement of absorbance at 560 nm and 690 nm. The concentration achieving 50% inhibition of cytopathic effect (effective concentration; EC_50_) was defined in GraphPad Prism version 8.4.3 (GraphPad Software) with a variable slope (four parameters). Non-treated cells were used as a control for 100% inhibition.

### Expression and purification of SARS-CoV-2 3CL^pro^

Expression plasmid (pGEX-5X-3-SARS-CoV-2-3CL) was a gift from Alejandro Chavez & David Ho & Sho Iketani (Addgene plasmid # 168457 ; http://n2t.net/addgene:168457 ; RRID:Addgene_168457). Expression and purification of SARS-CoV-2-3CLpro (3CLpro) was performed as previously described^43^ with a slight modification. Briefly, after initial purification of GST-fused 3CLpro with on-column cleavage of GST-tag by Factor Xa, the supernatant containing 3CLpro was dialyzed against 20 mM Tris-HCl, pH 7.5, and subjected to anion-exchange chromatography on a Resource Q column (Cytiva). The sample was eluted with a linear gradient of 0–0.5 M NaCl. Fractions containing 3CLpro was concentrated using a centrifugal filter (Amicon Ultra-15 centrifugal filter device with MWCO of 3 kDa) and applied onto a Superdex increase 200 10/300 column equilibrated with 20 mM Tris-HCl, pH 7.5, 150 mM NaCl, 1 mM EDTA for further purification by size exclusion chromatography. Collected fractions were analyzed by SDS-PAGE followed by Coomassie blue staining. Fractions containing 3CLpro were then dialyzed against 20 mM Tris-HCl, pH 7.5, 150 mM NaCl, 1 mM EDTA, and 1 mM TCEP, and concentrated to ∼10 mg/mL for crystal screening.

### Crystallization of SARS-CoV-2 3CL^pro^ with YH-6 (9a), data collection and processing

Crystals of 3C-like protease in complex with **8a** (**YH-6**) were obtained by the method of co-crystallization. Prior to the crystallization step, the complex of protein and **8a** (**YH-6**) was prepared at a protein concentration of 11 mg/mL and a protein : inhibitor molar ratio of 1:2. Crystals of this complex were grown by mixing with an equal volume of a solution composed of 16% (w:v) polyethylene glycol monomethyl ether, 20% glycerol, 80 mM TRIS (pH 8.5) and 8 mM NiCl_2_ using the sitting drop method. Suitable crystals appeared within one week, and were harvested and transferred to liquid nitrogen for storage until data collection. Diffraction data from a single crystal were collected in beamline BL5A at the Photon Factory (Tsukuba, Japan) under cryogenic conditions (100 K). Diffraction images were processed with the program MOSFLM and merged and scaled with the program SCALA or AIMLESS of the CCP4 suite. The structure of the WT protein was determined by the molecular replacement method using the coordinates of 3C-like protease in the unbound form (PDB entry code 6Y2E) with the program PHASER. The model was refined with the program REFMAC5 and built manually with COOT. Initial coordinates for the ligand were generated with the electronic Ligand Builder and Optimization Workbench (eLBOW) module included in the PHENIX suite. Validation was carried out with PROCHECK. Data collection and structure refinement statistics are given in Table S3. The coordinates and structure factors of 3C-Like protease in complex with the covalent reversible inhibitor **8a** (**YH-6**) have been deposited in the Protein Data Bank with entry code 7XAR.

### Animals and treatments

Seven-week-old male ICR/c mice (Kyudo Co., Ltd.) were housed under a standardized light-dark cycle at 24 ± 1°C and 60% ± 10% humidity with food and water ad libitum. A solution of YH6 was prepared by dissolving in dimethyl sulfoxide or 50% dimethyl sulfoxide, 20% ethanol,30% saline. The drugs were administrated orally using metal gavage needles or intravenously using a 30-gauge needle. All experimental procedures were performed under the approval and guidelines of Kyushu University.

### Evaluation of 8a (YH-6) concentrations in mouse serum and lung

The protein precipitation solvent (79.5% acetonitrile, 20% methanol, and 0.5% trichloroacetic acid) was added to the plasma samples in a ratio of 50/50 (v/v). Then, plasma added the protein precipitation solvents was separated by centrifugation (12000 rpm, 20 min). The supernatant was used as the sample for LC/MS/MS. Lung from mouse were collected and, excised lungs were put in 5 ml ice cold saline and cut into 20 mg pieces. Water 200 µL was added to the lung pieces samples and, samples were homogenized using handy homogenizer. Then, the lysate added the protein precipitation solvents was separated by centrifugation (12000 rpm, 20 min). The supernatant was used as the sample for LC/MS/MS. Drugs were resolved using an AQUITY UPLC HSS PFP column (50 mm × 2.1 mm, 50 mm, p/n186005965, Waters, Milford, MA, USA) on an AQUITY UPLC H-Class XHCLQT0100 system (Waters), consisting of a vacuum degasser, binary pump, thermostatically controlled column compartment, thermostatically controlled autosampler, and a diode array detector. Mobile-phase A consisted of 0.1% formic acid/water; mobile-phase B consisted of 0.1% formic acid/acetonitrile. A linear gradient was generated at 1 ml/min: 0–0.1 min, 80% A, and 20% B; 0.1–2.8 min, 20% A, and 80% B; 2.8–6.0 min, 80% A, and 20% A (Waters application note). The injection volume was 10 µL. The column temperature was controlled at 40 °C and the autosampler compartment was set to 10 °C. An AQUITY TQ mass spectrometer controlled by MasslynxTM software (Waters) was operated in selected reaction monitoring mode. The multiple reaction monitor was set at a mass-to-charge ratio (m/z) of 498.1–161.9 m/z (cone voltage:20, collision voltage:30) for YH6, 493–117 m/z (cone voltage: 30, collision voltage: 20) for internal standard (cilnidipine) (Car no. 036-25101, Fujifilm).

## Supporting Information

**Figure S1.**
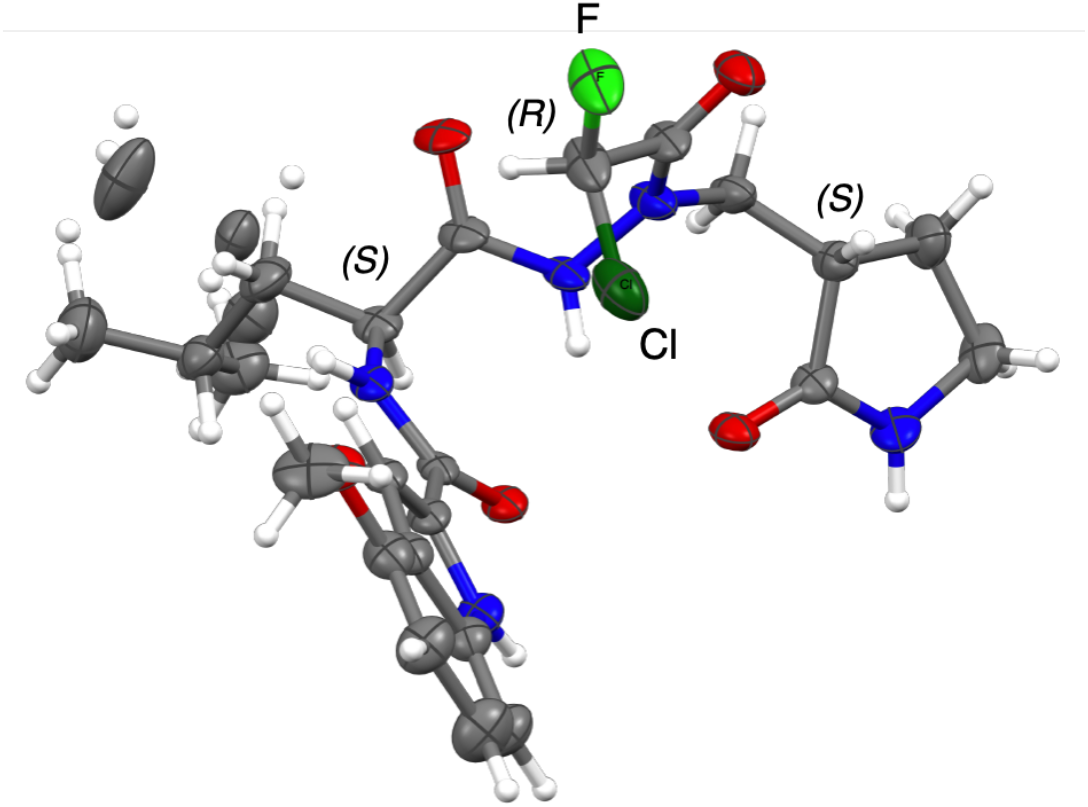
X-ray crystallographic structure of (*S, S, R*) -**8b**. The crystal data and structure refinement statistics were shown in Table S1.

**Figure S2.**
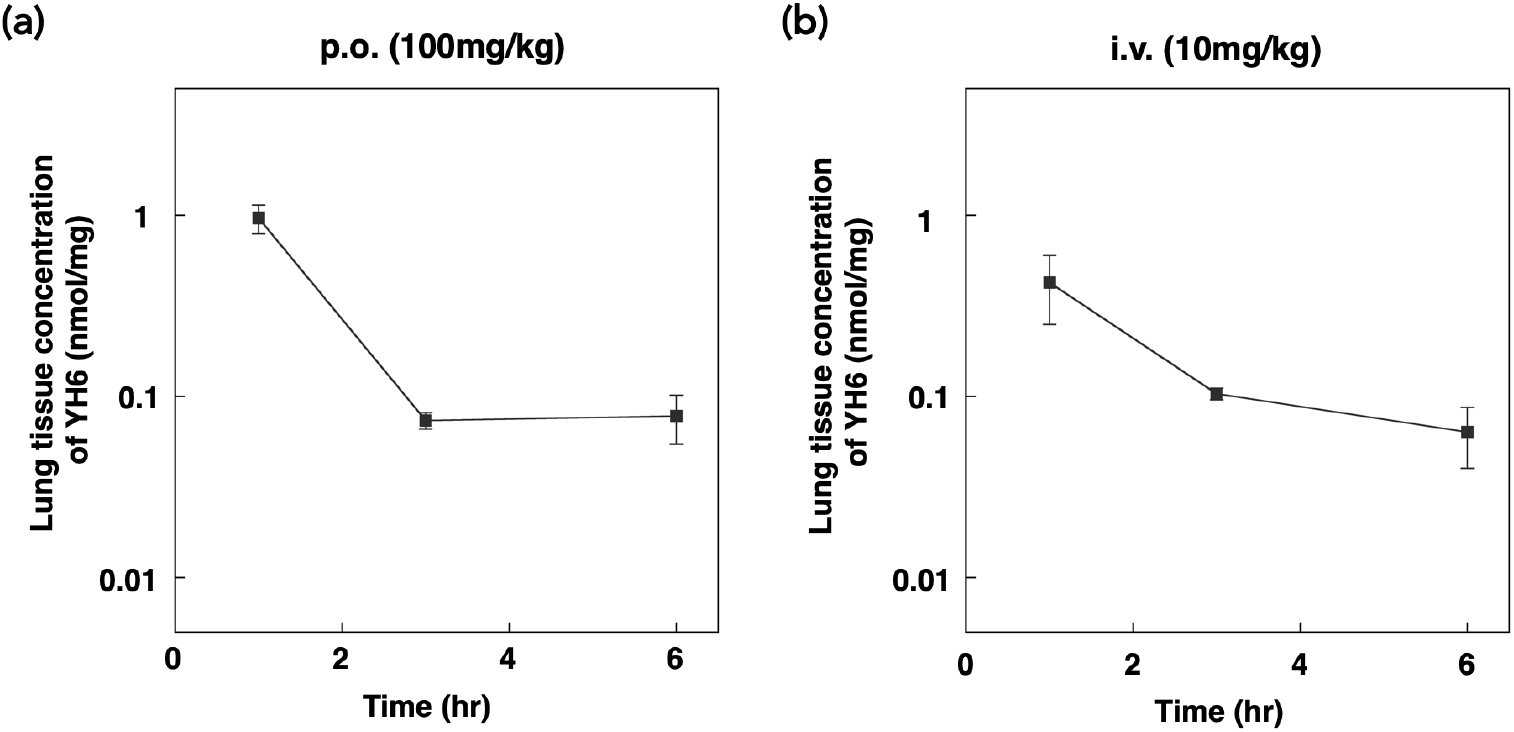
Mean lung tissue concentration (nmol/mg)-time profile of **8a** (**YH-6**) in mice (fasted state) after (a) p.o. (100 mg/kg) and (b) i.v. (10 mg/kg) administration. Data are shown as the mean ± SE (n = 3).

**Table S1.** Crystal data and structure refinement statistics of **8b**.

|  |  |
| --- | --- |
| Empirical formula | C<sub>23</sub>H<sub>29</sub>ClFN<sub>5</sub>O<sub>5</sub> |
| Formula weight | 509.96 |
| Temperature/K | 100.00(10) |
| Crystal system | hexagonal |
| Space group | P6<sub>1</sub> |
| a/Å | 23.7450(4) |
| b/Å | 23.7450(4) |
| c/Å | 10.0317(2) |
| α/° | 90 |
| β/° | 90 |
| γ/° | 120 |
| Volume/Å<sup>3</sup> | 4898.4(2) |
| Z | 6 |
| ρ<sub>calc</sub>/g/cm<sup>3</sup> | 1.037 |
| μ/mm<sup>-1</sup> | 1.373 |
| F(000) | 1608.0 |
| Crystal size/mm<sup>3</sup> | 0.209 × 0.049 × 0.047 |
| Radiation | Cu Kα (λ = 1.54184) |
| 2θ range for data collection/° | 7.446 to 151.7 |
| Index ranges | -29 ≤ h ≤ 29, -29 ≤ k ≤ 29, -9 ≤ l ≤ 11 |
| Reflections collected | 83295 |
| Independent reflections | 6045 [R<sub>int</sub> = 0.0338, R<sub>sigma</sub> = 0.0136] |
| Data/restraints/parameters | 6045/7/349 |
| Goodness-of-fit on F<sup>2</sup> | 1.111 |
| Final R indexes [I ≥ 2σ (I)] | R<sub>1</sub> = 0.0473, wR<sub>2</sub> = 0.1326 |
| Final R indexes [all data] | R<sub>1</sub> = 0.0478, wR<sub>2</sub> = 0.1331 |
| Largest diff. peak/hole / e Å<sup>-3</sup> | 0.32/-0.25 |
| Flack parameter | 0.04(2) |

**Table S2.** Antiviral activity against various SARS-CoV-2 strains in cytopathic effect inhibition assay with 293TAT cell.

| | 293TAT $\alpha$ EC<sub>50</sub> (nM) | | | 293TAT $\delta$ EC<sub>50</sub> (nM) | | |
| --- | --- | --- | --- | --- | --- | --- |
|  | HSA 0% | HSA 1% | HSA 2% | HSA 0% | HSA 1% | HSA 2% |
| **8a (YH-6)** | 7.1 | 18.7 | 24.4 | 12.4 | 39.1 | 57.9 |
| **8b** | 9.1 | 13 | 16.1 | 21.3 | 25.7 | 34.4 |
| NHC | 118.8 | 353.8 | 336.9 | 492.8 | 497.1 | 465.6 |

**Table S3.** Data collection and refinement statistics of X-ray crystal structure of SARS-CoV-2 M^pro^ in complex with **8a** (**YH-6**).

|  |  |
| --- | --- |
| Data Collection |  |
| Space Group | P 1 2<sub>1</sub> 1 |
| Unit cell |  |
| a, b, c (Å) | 47.0, 63.4, 102.8 |
| α, β, γ (°) | 90.0, 90.4 90.0 |
| Resolution (Å) | 47.0 - 1.60 |
| Wavelength | 1.0000 |
| Observations | 542,961 (27,343) |
| Unique reflections | 79,374 (3,907) |
| *R*<sub>merge</sub> | 0.059 (0.883) |
| *R*<sub>p.i.m.</sub> | 0.024 (0.356) |
| CC<sub>1/2</sub> | 0.999 (0.815) |
| *I* / σ (*I*) | 18.3 (2.2) |
| Multiplicity | 6.8 (7.0) |
| Completeness (%) | 99.5 (98.6) |
| Refinement Statistics |  |
| Resolution (Å) | 47.0 - 1.60 |
| *R*<sub>work</sub> / *R*<sub>free</sub> (%) | 17.3 / 20.7 |
| No. protein chains | 2 |
| No. atoms |  |
| Protein | 4,788 |
| Ligand | 66 |
| Other (no solvent) | 8 |
| Water | 473 |
| B-factor (Å<sup>2</sup>) |  |
| Protein | 24.1 |
| Ligand | 23.7 |
| Other (no solvent) | 39.9 |
| Water | 30.7 |
| Ramachandran Plot |  |
| Preferred (%) | 91.5 |
| Allowed (%) | 8.2 |
| Outliers (%) | 0.4 |
| RMSD Bond (Å) | 0.009 |
| RMSD Angle (°) | 1.55 |
| PDB entry code | 7XAR |

Statistical values given in parenthesis refer to the highest resolution bin.

### General synthetic methods

Reagents and solvents were obtained from commercial suppliers and used without further purification, unless otherwise stated. Reactions were carried out under a positive atmosphere of nitrogen, unless otherwise stated. Reactions were monitored by thin layer chromatography (TLC) carried out on Merck TLC Silica gel 60 F_254_, using shortwave UV light as the visualizing agent and phosphomolybdic acid in EtOH and heat as developing agent. Flash column chromatography was performed using Kanto Chemical Silica gel 60 N (spherical, 40-50 µm) or using Biotage^®^ Isolera medium pressure column chromatography system and Sfär Silica HC D 20 µm. ^1^H and ^13^C NMR spectra were recorded on Bruker Avance III HD 500 MHz spectrometer and were calibrated using residual undeuterated solvent as the internal references (CDCl_3_: 7.26 ppm; CD_3_OD: 3.31 ppm; DMSO-*d*_6_: 2.50 ppm). The following abbreviations were used to explain NMR peak multiplicities: s = singlet, d = doublet, t = triplet, q = quartet, p = pentet, m = multiplet, br = broad. Low-resolution and high-resolution mass spectra were recorded on Bruker micrOTOF focus II mass spectrometer using electrospray ionization time-of-flight (ESI-TOF) reflectron experiments. X-ray crystallographic experiments were performed using Rigaku FR-E+ instrument. S18

### Benzyl 3-formyl-2-oxopyrrolidine-1-carboxylate (2)

To a solution of **1** (13.2 g, 60.0 mmol) in toluene (60 mL) was added Bredereck’s reagent (13.5 mL, 66.0 mmol) under argon. The mixture was refluxed for 1h, then cooled to room temperature, and the reaction mixture was concentrated under reduced pressure. The residue was washed with hexane, dried *in vacuo* to give slightly yellow solid. To a solution of this slightly yellow solid in THF (300 mL) was added 1M HCl (90 mL). The reaction mixture was stirred for 1 h at room temperature, then poured into satturated NaHCO_3_ and adjust pH to 7.0. The aqueous layer was extracted with ethyl acetate, washed with saturated brine, dried over MgSO_4_, filtrated, and concentrated *in vacuo*. The residue was purified by flush silica gel column chromatography to give **2** (15.5 g, quant.) as yellow oil.

^1^H NMR (500 MHz, DMSO-*d*_6_): ’ 7.96 (1H, br), 7.31-7.44 (5H, m), 7.00 (1H, t, *J* = 1.8 Hz), 5.19 (2H, s), 3.63 (2H, t, *J* = 7.5 Hz), 2.89-2.92 (2H, m). HRMS (ESI): *m/z* [M-H]^-^ calcd for C_13_H_12_NO_4_ 246.0772; found 246.0759.

### Benzyl 3-((2-(((benzyloxy)carbonyl)-L-leucyl)hydrazineyl)methyl)-2-oxopyrrolidine-1-carboxylate (4)

To a solution of **2** (10.9 g, cal. 41.9 mmol) in EtOH (28 mL) was added **3**^s1^ (11.7 g, 41.9 mmol) under argon. The mixture was refluxed for 30 min, then cooled to room temperature, concentrated under reduced pressure to remove the solvent. To a solution of this residue in CH_2_Cl_2_ (80 mL) was added dimethylamine-borane complex (3.95 g, 67.1 mmol) and cooled to 0 °C.^s2^ Then a solution of p-toluene sulfonic acid monohydride (4.50 g, 237 mmol) in CH_2_Cl_2_/MeOH (3:1, 80 mL) was added. After stirred for 30 min at 0 °C, 10% aqueous Na_2_CO_3_ (240 mL) and MeOH (80 mL) were added, and the mixture refluxed for 30 min. Then organic layer was separated and aqueous layer was extracted with CH_2_Cl_2_. The organic layer was combined and washed with saturated brine, dried over MgSO_4_, filtrated, and concentrated *in vacuo*. The residue was purified by flush silica gel column chromatography to give **4** (14.5 g, 68%) as a colorless solid.

^1^H NMR (500 MHz, DMSO-*d*_6_): ’ 9.43 and 9.45 (1H, d, *J* = 6.5 Hz), 7.29-7.43 (10H, m), 5.21 (2H, s), 5.07 (1H, m), 5.01 (2H, s), 3.95 (1H, m), 3.73 (1H, m), 3.58 (1H, m), 3.02-3.07 (1H, m), 2.66-2.74 (2H, m), 2.08-2.29 (1H, m), 1.77-1.83 (1H, m), 1.49-1.60 (1H, m), 1.43-1.48 (1H, m), 1.19 -1.34 (1H, m), 0.86 (3H, d, *J* = 6.5 Hz), 0.84 (3H, d, *J* = 6.5 Hz). ^13^C-NMR (125 MHz, DMSO-*d*_6_): δ 174.5, 171.1 and 171.2, 155.8, 150.9, 137.1, 135.8, 128.4, 128.3, 128.1, 127.7, 127.6, 66.9, 65.3, 51.7, 51.3 and 51.4, 44.4, 42.2 and 42.3, 24.2, 22.8, 22.5 and 22.6, 21.6. HRMS (ESI): *m/z* [M+H]^+^ calcd for C_27_H_34_N_4_NaO_6_ 533.2371; found 533.2386.

### Benzyl 3-((2-(((benzyloxy)carbonyl)-L-leucyl)-1-(*tert*-butoxycarbonyl)hydrazineyl)methyl)-2-oxopyrrolidine-1-carboxylate (5)

To a solution of **4** (14.5 g, 28.3 mmol) in CH_2_ Cl_2_ (90 mL) was added Boc_2_O (6.54 g, 30.0 mmol) and DMAP (299 mg, 2.45 mmol) under argon. The mixture was stirred for 1 h at room temperature, then poured into saturated NH_4_Cl. The aqueous layer was extracted with CH_2_Cl_2_, and the organic layer was combined, and washed with saturated brine, dried over MgSO_4_, filtrated, and concentrated *in vacuo*. The residue was purified by flush silica gel column chromatography to give **5** (13.6 g, 79%) as a colorless solid.

^1^H NMR (500 MHz, DMSO-*d*_6_): ’ 10.10 and 10.15 (1H, br), 7.26-7.48 (10H, m), 5.18-5.24 (2H, m), 4.98-5.06 (2H, m), 4.01-4.05 (1H, m), 3.42-3.72 (4H, m), 2.72-2.81 (1H, m), 2.18 (1H, m), 1.74-1.80 (1H, m), 1.60-1.68 (1H, m), 1.40-1.52 (2H, m), 1.33 (9H, s), 0.84-0.87 (6H, m). ^13^C-NMR (125 MHz, DMSO-*d*_6_): δ 173.5, 171.2, 155.9, 154.8, 150.9, 137.0 and 137.1, 135.8, 128.4, 128.2 and 128.3, 128.0, 127.7, 127.6, 80.1, 66.9, 65.3, 51.4, 44.2, 42.2, 27.7, 24.1, 22.9, 22.5, 21.3 and 21.4, 20.7. HRMS (ESI): *m/z* [M+Na]^+^ calcd for C_32_H_42_N_4_NaO_8_ 633.2895; found 633.2450.

### *tert*-Butyl 2-(L-leucyl)-1-((2-oxopyrrolidin-3-yl)methyl)hydrazine-1-carboxylate (6)

A suspension of **5** (3.00 g, 4.91 mmol) and palladium-activated carbon (Pd 10%, 450 mg) in methanol (40 mL) was vigorously stirred for 1 h under hydrogen before inside air was replaced with hydrogen. Then palladium-activated carbon (Pd 10%, 300 mg) in methanol (10 mL) was added and stirred for 1 h. The reaction mixture was filtrated through celite pad with methanol, and the filtrate was concentration *in vacuo*. The residue was purified by medium pressure column chromatography (Biotage^®^ Sfär Silica HC D 20 µm 25 g connected with 10 g, chloroform:methanol = 100:0 to 70:30) to give diastereomeric mixture of **6** (1.60 g, 95%) as a colorless amorphous powder.

### *tert*-Butyl 2-(L-leucyl)-1-(((*S*)-2-oxopyrrolidin-3-yl)methyl)hydrazine-1-carboxylate ((*S,S*)-6) and *tert*-Butyl 2-(L-leucyl)-1-(((*R*)-2-oxopyrrolidin-3-yl)methyl)hydrazine-1-carboxylate ((*S,R*)-6)

The disasereomeric mixture of **6** was separation by medium pressure chiral column chromatography (Daicel CHIRALFLASH^®^ IC, 20 µm 30β x 100 mm 40 g, hexane : ethanol : diethylamine = 50 : 50 : 0.05, flow rate: 12 mL/min, detector: UV 200/204 nm, repeated 20 times) to give **(*S***,***S*)-6** (576 mg, less polar fraction) and **(*S***,***R*)-6** (614 mg, more polar fraction) as a colorless amorphous powder, respectively.

**(*S***,***S*)-6**: ^1^H-NMR (500 MHz, DMSO-*d*_6_): δ 9.97 (1H, br), 7.65 (1H, br), 3.12-3.54 (4H, m), 2.44 (1H, m), 2.22 (1H, m), 1.71-1.81 (2H, m), 1,25-1.44 (2H, m), 1.36 (9H, s), 0.88 (3H, d, *J* = 6.5 Hz), 0.85 (3H, d, *J* = 7.0 Hz). ^13^C-NMR (125 MHz, DMSO-*d*_6_): δ 176.7, 173.7, 154.8, 79.8, 51.5, 49.5, 43.5, 43.4, 41.4, 27.8, 26.0, 23.9, 23.8, 23.0, 21.9. HRMS (ESI): *m/z* [M+H]^+^ calcd for C_16_H_31_N_4_O_4_ 343.2340; found 343.2309.

**(*S***,***R*)-6**: ^1^H-NMR (500 MHz, DMSO-*d*_6_): δ 9.98 (1H, br), 7.66 (1H, br), 3.42-3.54 (2H, m), 3.11-3.24 (2H, m), 2.43-2.48 (1H, m), 2.20-2.25 (1H, m), 1.72-1.80 (2H, m), 1,25-1.44 (2H, m), 1.35 (9H, s), 0.89 (3H, d, *J* = 6.5 Hz), 0.86 (3H, d, *J* = 7.0 Hz). ^13^C-NMR (125 MHz, DMSO-*d*_6_): δ 176.6, 173.7, 154.9, 79.8, 51.5, 49.5, 43.7, 43.4, 41.4, 27.8, 26.0, 23.9, 23.8, 23.0, 21.6. HRMS (ESI): *m/z* [M+H]^+^ calcd for C_16_H_31_N_4_O_4_ 343.2340; found 343.2302.

### *tert*-Butyl 2-((4-fluoro-1*H*-indole-2-carbonyl)-L-leucyl)-1-(((*S*)-2-oxopyrrolidin-3-yl)methyl)hydrazine-1-carboxylate (7a)

To a solution of 4-fluoroindole-2-carboxylic acid (212 mg, 1.16 mmol) in DMF (3 mL) was added COMU (586 mg, 1.34 mmol) and TEA (374 µL, 2.68 mmol) at 0 °C under argon. The mixture was stirred for 15 min at room temperature, then re-cooled to 0 °C, and **(*S***,***S*)-6** (307 mg, 0.890 mmol) in DMF (1 mL) was added. The reaction mixture was stirred for 30 min at room temperature, then poured into ethyl acetate and saturated aqueous NaHCO_3_. The organic layer was separated, washed with saturated brine, dried over MgSO_4_, filtrated, and concentrated *in vacuo*. The residue was purified by medium pressure column chromatography (Biotage^®^ Sfär Silica HC D 20 µm 10 g, chloroform:methanol = 100:0 to 96:4) to give **7a** (357 mg, 80%) as a yellow solid.

^1^H-NMR (500 MHz, DMSO-*d*_6_): δ 11.91 (1H, br), 10.21 (1H, br), 8.60 (1H, br), 7.26 (1H, d, *J* = 8.5 Hz), 7.15 (1H, ddd, *J* = 13.5, 8.8, 5.5 Hz), 6.81 (1H, dd, *J* = 10.5, 8.0 Hz), 4.55 (1H, br), 3.57 (1H, m), 3.37 (1H, m), 3.08-3.17 (2H, m), 2.42-2.45 (1H, m), 2.23-2.25 (1H, m), 1.70-1.80 (3H, m), 1.54-1.59 (1H, m), 1.34 (9H, s), 0.93 (3H, d, *J* = 6.5 Hz), 0.90 (3H, d, *J* = 6.5 Hz). ^13^C-NMR (125 MHz, DMSO-*d*_6_): δ 176.5, 170.9, 160.5, 156.1 (d, *J*_(C–F)_ = 244.3 Hz), 154.8, 138.9 (d, *J*_(C–F)_ = 10.6 Hz), 131.6, 123.9 (d, *J*_(C–F)_ = 7.6 Hz), 116.2 (d, *J*_(C–F)_ = 21.9 Hz), 108.9, 103.9 (d, *J*_(C–F)_ = 18.4 Hz), 99.0, 79.8, 50.0. 49.8, 27.8, 26.5, 24,3, 23.0, 21.3. HRMS (ESI): *m/z* [M+Na]^+^ calcd for C_25_H_34_FN_5_NaO_5_ 526.2436; found 526.2244.

### *tert*-Butyl 2-((4-methoxy-1*H*-indole-2-carbonyl)-L-leucyl)-1-(((*S*)-2-oxopyrrolidin-3-yl)methyl)hydrazine-1-carboxylate (7b)

^1^H-NMR (500 MHz, CDCl_3_): δ 11.55 (1H, br), 10.16 (1H, br), 8.42 (1H, br), 7.64 (1H, br), 7.36 (1H, d, *J* = 1.5 Hz), 7.09 (1H, dd, *J* = 8.0, 8.0 Hz), 7.00 (1H, d, *J* = 8.0 Hz), 6.50 (1H, d, *J* = 7.5 Hz), 4.52 (1H, br), 3.88 (3H, s), 3.52-3.62 (1H, m), 2.27-3.44 (1H, m), 3.05-3.08 (2H, m), 2.39-2.46 (1H, m), 2.20-2.27 (1H, m), 1.67-1.83 (3H, m), 1.51-1.58 (1H, m), 1.34 (1H, s), 0.93 (3H, d, *J* = 6.5 Hz), 0.89 (3H, d, *J* = 6.5 Hz),

^13^C-NMR (125 MHz, CDCl_3_): δ 180.1 and 179.5, 171.4 and 170.5, 162.7, 154.8, 154.2, 138.9, 128.8, 125.9, 118.8, 106.4, 100.4, 99.5, 81.7 and 81.3, 63.2, 55.4, 51.8, 41.1, 40.7, 40.5, 28.4 and 28.3, 25.1, 23.9 and 23.7, 23.4, 22.4 and 21.2. HRMS (ESI): *m/z* [M+Na]^+^ calcd for C_26_H_37_N_5_NaO_6_ 538.2636; found 538.2570

### *N*-((*S*)-1-(2-((*R*)-2-Chloro-2-fluoroacetyl)-2-(((*S*)-2-oxopyrrolidin-3-yl)methyl)hydrazineyl)-4-methyl-1-oxopentan-2-yl)-4-fluoro-1*H*-indole-2-carboxamide (8a, YH-6)

A solution of **7a** (328 mg, 0.651 mmol), HCl/dioxane (4 M, 6.5 mL) was stirred for 50 min atroom temperature under argon. The reaction mixture was concentrated in vacuo, and the residue was dissolved to DMF (6 mL), then (*R*)-CFA (45.7% in 1,4-dioxane, 140 µL, 0.722 mmol), TEA (453 µL, 3.25 mmol), and COMU (429 mg, 1.00 mmol) was added at 0 °C. The reaction mixture was stirred at room temperature for 10 min, followed by adding COMU (284 mg, 0.663 mmol and 57 mg, 0.133 mmol). After 20 min, the reaction mixture was poured into saturated aqueous NaHCO_3_ and extracted with ethyl acetate. The organic layer was separated, washed with saturated brine, dried over MgSO_4_, filtrated, and concentrated *in vacuo*. The residue was purified by medium pressure column chromatography (Biotage^®^ Sfär Silica HC D 20 µm 10 g, chloroform:methanol = 100:0 to 96:4) to give **8a** (205 mg, 63%) as a yellow solid.

^1^H-NMR (500 MHz, DMSO-*d*_6_): δ 11.94 (1H, br), 11.11 (1H, br), 8.82 (1H, br), 7.43 (1H, br), 7.28 (1H, d, *J* = 8.0 Hz), 7.17 (1H, ddd, *J* = 13.5, 8.0, 5.5 Hz), 6.90 (1H, d, *J* = 50.5 Hz), 6.81 (1H, dd, *J* = 10.5, 8.0 Hz), 4.34 (1H, br), 3.91 (1H, m), 3.13-3.26 (2H, m), 2.54-2.61 (1H, m), 2.15-2.20 (1H, m), 1.82-1.83 (3H, m), 1.77-1.80 (1H, m), 0.98 (3H, d, *J* = 6.5 Hz), 0.93 (3H, d, *J* = 6.0 Hz). ^13^C-NMR (125 MHz, DMSO-*d*_6_): δ 175.9, 172.6, 165.9 (d, *J*_(C–F)_ = 25.0 Hz), 161.6, 156.2 (d, *J*_(C–F)_ = 245.0 Hz), 139.0 (d, *J*_(C–F)_ = 10.0 Hz), 131.0, 124.2 (d, *J*_(C–F)_ = 7.5 Hz), 116.2 (d, *J*_(C–F)_ = 21.3 Hz), 109.0, 104.0 (d, *J*_(C–F)_ = 18.8 Hz), 90.1(d, *J*_(C–F)_ = 241.3 Hz), 65.9, 50.8. 48.7, 38.7, 25.7, 24.3, 23.0, 21.3. HRMS (ESI): *m/z* [M+Na]^+^ calcd for C_22_H_26_ClF_2_N_5_NaO_4_ 520.1534; found 520.1351.

### *N*-((*S*)-1-(2-((*R*)-2-Chloro-2-fluoroacetyl)-2-(((*S*)-2-oxopyrrolidin-3-yl)methyl)hydrazineyl)-4-methyl-1-oxopentan-2-yl)-4-methoxy-1*H*-indole-2-carboxamide (8b)

^1^H-NMR (500 MHz, CDCl_3_): δ 11.60 (1H, br), 10.56 (1H, br), 7.43 (1H, br), 7.25 (1H, dd, *J* = 8.5, 7.0 Hz), 7.18 (1H, d, *J* = 8.5 Hz), 7.12 (1H, d, *J* = 2.0 Hz), 6.51 (1H, d, *J* = 7.5 Hz), 6.31 (1H, d, *J* = 50.0 Hz), 6.33 (1H, d, *J* = 7.5 Hz), 4.97 (1H, br), 4.07-4.11 (1H, m), 2.65-2.70 (2H, m), 2.39 (1H, m), 2.09-2.40 (2H, m), 1.62-1.85 (4H, m), 1.06 (3H, d, *J* = 7.0 Hz), 1.02 (3H, d, *J* = 6.0 Hz). ^13^C-NMR (125 MHz, CDCl_3_): δ 179.8, 173.0, 164.3 (d, *J*_(C–F)_ = 25.3 Hz), 162.8, 139.1, 128.2, 126.4, 118.8, 106.3, 100.4, 90.3 (d, *J*_(C–F)_ = 247.0 Hz), 55.5, 52.4. 49.3, 25.3, 23.7, 23.4, 21.7. HRMS (ESI): *m/z* [M+Na]^+^ calcd for C_23_H_29_ClFN_5_NaO_5_ 532.1733; found 532.1767.

### *N*-((*S*)-1-(2-((*S*)-2-Chloro-2-fluoroacetyl)-2-(((*S*)-2-oxopyrrolidin-3-yl)methyl)hydrazineyl)-4-methyl-1-oxopentan-2-yl)-4-fluoro-1*H*-indole-2-carboxamide (9)

^1^H-NMR (500 MHz, DMSO-*d*_6_): δ 11.96 (1H, s), 10.7 and 10.83 (1H, s), 8.81 and 8.74 (1H, d, *J* = 7.0 Hz and 6.0 Hz), 7.74 and 7.78 (1H, s), 7.42 (1H, br-d), 7.28 (1H, br-d), 7.16-7.19 (1H, m), 6.61 and 7.05 (1H, d, *J* = 49.5 Hz), 4.47 (1H, m), 3.91 (1H, m), 3.75 and 3.87 (1H, m), 3.51 and 3.56 (1H, m), 3.29-3.37 (1H, m), 3.05-3.20 (2H, m), 2.52-2.57 (1H, m), 2.20 (1H, m), 1.57-1.87 (4H, m), 0.97 (3H, m), 0.92 (3H, m). ^13^C-NMR (125 MHz, DMSO-*d*_6_): δ 176.1, 170.9 and 172.7, 164.6 and 164.9 (d, *J*_(C–F)_ = 26.3 and 25.0 Hz), 161.2 and 161.4, 156.2 (d, *J*_(C–F)_ = 243.8 Hz), 139.0 (d, *J*_(C–F)_ = 10.0 Hz), 131.2 and 131.3, 124.1 (d, *J*_(C–F)_ = 7.5 Hz), 116.2 (d, *J*_(C–F)_ = 21.3 Hz), 109.0, 104.0 (d, *J*_(C–F)_ = 17.5 Hz), 99.3 and 99.4, 90.6 and 90.9 (d, *J*_(C–F)_ = 241.3 and 240.0 Hz), 65.9, 51.3. 50.4, 49.6, 48.3, 47.0, 39.0, 26.1 and 26.6, 24.3, 21.2 and 21.5, 21.3. HRMS (ESI): *m/z* [M+Na]^+^ calcd for C_22_H_26_ClF_2_N_5_NaO_4_ 520.1534; found 520.1528.

### *tert*-Butyl 2-((4-fluoro-1*H*-indole-2-carbonyl)-L-leucyl)-1-(((*R*)-2-oxopyrrolidin-3-yl)methyl)hydrazine-1-carboxylate (13a)

To a solution of 4-fluoroindole-2-carboxylic acid (68 mg, 0.380 mmol) in DMF (2 mL) was added COMU (188 mg, 0.439 mmol) and TEA (120 µL, 0.861 mmol) at 0 °C under argon. The mixture was stirred for 15 min at room temperature, then re-cooled to 0 °C, and **(*S***,***R*)-6** (98 mg, 0.286 mmol) in DMF (1 mL) was added. The reaction mixture was stirred for 30 min at room temperature, then poured into ethyl acetate and saturated aqueous NaHCO3. The organic layer was separated, washed with saturated brine, dried over MgSO4, filtrated, and concentrated in vacuo. The residue was purified by medium pressure column chromatography (Biotage® Sfär Silica HC D 20 µm 10 g, chloroform:methanol = 100:0 to 90:10) to give **13a** (114 mg, 81%) as a yellow solid.

^1^H-NMR (500 MHz, DMSO-*d*_6_): δ 11.91 (1H, br), 10.20 (1H, br), 8.63 (1H, br), 7.66 (1H, s), 7.40 (1H, d, *J* = 1.5 Hz), 7.26 (1H, d, *J* = 8.5 Hz), 7.15 (1H, ddd, *J* = 13.5, 8.0, 5.5 Hz), 6.81 (1H, dd, *J* = 10.5, 8.0 Hz), 4.55 (1H, br), 3.40-3.55 (2H, m), 3.11-3.19 (2H, m), 2.46 (1H, m), 2.23 (1H, m), 1.73-1.79 (3H, m), 1.57-1.60 (1H, m), 1.35 (9H, s), 0.93 (3H, d, *J* = 6.5 Hz), 0.90 (3H, d, *J* = 6.5 Hz). ^13^C-NMR (125 MHz, DMSO-*d*_6_): δ 176.5, 171.0, 160.5, 156.2 (d, *J*_(C–F)_ = 244.3 Hz), 154.8, 138.9 (d, *J*_(C–F)_ = 10.8 Hz), 131.6, 123.9 (d, *J*_(C–F)_ = 7.5 Hz), 116.2 (d, *J*_(C–F)_ = 21.8 Hz), 108.8, 103.9 (d, *J*_(C–F)_ = 18.0 Hz), 99.0, 79.9, 49.8, 27.8, 26.3, 24,3, 23.0, 21.3. HRMS (ESI): *m/z* [M+Na]^+^ calcd for C_25_H_34_FN_5_NaO_5_ 526.2436; found 526.2282.

### *N*-((*S*)-1-(2-((*R*)-2-chloro-2-fluoroacetyl)-2-(((*R*)-2-oxopyrrolidin-3-yl)methyl)hydrazineyl)-4-methyl-1-oxopentan-2-yl)-4-fluoro-1*H*-indole-2-carboxamide (10a)

A solution of **13a** (100 mg, 0.199 mmol), HCl/dioxane (4 M, 2.0 mL) was stirred for 50 min at room temperature under argon. The reaction mixture was concentrated in vacuo, and the residue was dissolved to DMF (3 mL), then (*R*)-CFA (45.7% in 1,4-dioxane, 47 µL, 0.242 mmol), TEA (238 µL, 1.70 mmol), and COMU (131 mg, 0.306 mmol) was added at 0 °C. The reaction mixture was stirred at room temperature for 10 min, followed by adding COMU (88 mg, 0.205 mmol, 44 mg, 0.103 mmol, and 44 mg, 0.103 mmol) and TEA (95 µL, 0.681 mmol). After 30 min, the reaction mixture was poured into saturated aqueous NaHCO_3_ and extracted with ethyl acetate. The organic layer was separated, washed with saturated brine, dried over MgSO_4_, filtrated, and concentrated *in vacuo*. The residue was purified by medium pressure column chromatography (Biotage^®^ Sfär Silica HC D 20 µm 10 g, chloroform:methanol = 100:0 to 95:5) to give **10a** (26 mg, 26%) as a yellow solid.

^1^H-NMR (500 MHz, DMSO-*d*_6_): δ 11.94 (1H, br), 11.04 (1H, br), 7.43 (1H, br), 8.78 (1H, d, *J* = 6.5 Hz), 7.73 (1H, br), 7.42 (1H, br), 7.28 (1H, d, *J* = 8.0 Hz), 7.17 (1H, ddd, *J* = 13.0, 8.0, 5.5 Hz), 6.96 (1H, d, *J* = 52.0 Hz), 6.82 (1H, dd, *J* = 10.5, 8.0 Hz), 4.35 and 4.50 (1H, br), 3.85 (1H, m), 3.41 (1H, m), 3.11-3.21 (2H, m), 2.43 and 2.60 (1H, m), 2.23 (1H, m), 1.52-1.90 (4H, m), 0.98 (3H, d, *J* = 6.0 Hz), 0.93 (3H, d, *J* = 6.0 Hz). ^13^C-NMR (125 MHz, DMSO-*d*_6_): δ 175.8, 172.9, 165.6 (d, *J*_(C–F)_ = 26.7 Hz), 161.7, 156.1 (d, *J*_(C–F)_ = 244.6 Hz), 139.0 (d, *J*_(C–F)_ = 10.7 Hz), 131.0, 124.2 (d, *J*_(C–F)_ = 6.9 Hz), 116.1 (d, *J*_(C–F)_ = 21.9 Hz), 109.0, 104.0 (d, *J*_(C–F)_ = 18.2 Hz), 99.4, 90.3 (d, *J*_(C–F)_ = 241.6 Hz), 65.9, 50.8, 49.4, 38.6, 26.2, 24.4, 23.2, 21.1. HRMS (ESI): *m/z* [M+Na]^+^ calcd for C_22_H_26_ClF_2_N_5_NaO_4_ 520.1534; found 520.1343.

### *tert*-Butyl 2-((4-methoxy-1*H*-indole-2-carbonyl)-L-leucyl)-1-(((*R*)-2-oxopyrrolidin-3-yl)methyl)hydrazine-1-carboxylate (13b)

To a solution of 4-methoxy-2-carboxylic acid (98.4 mg, 0.515 mmol) in DMF (2 mL) was added COMU (254.0 mg, 0.593 mmol) and TEA (162 µL, 1.16 mmol) at 0 °C under argon. The mixture was stirred for 15 min at room temperature, then re-cooled to 0 °C, and **(*S***,***R*)-6** (133 mg, 0.388 mmol) in DMF (2 mL) was added. The reaction mixture was stirred for 30 min at room temperature, then poured into ethyl acetate and saturated aqueous NaHCO_3_. The organic layer was separated, washed with saturated brine, dried over MgSO_4_, filtrated, and concentrated in vacuo. The residue was purified by medium pressure column chromatography (Biotage® Sfär Silica HC D 20 µm 5 g, chloroform:methanol = 100:0 to 90:10) to give **13b** (198 mg, 99%) as a light brown solid.

^1^H-NMR (500 MHz, DMSO-*d*_6_): δ 11.56 (1H, s), 10.16 (1H, br), 8.45 (1H, br), 7.66 (1H, s), 7.37 (1H, d, *J* = 2.0 Hz), 7.09 (1H, dd, *J* = 8.0, 7.5 Hz), 7.00 (1H, d, *J* = 8.0 Hz), 6.50 (1H, d, *J* = 7.5 Hz), 4.52 (1H, br), 3.88 (3H, s), 3.51 (1H, m), 3.40-3.45 (1H, m), 3.09-3.19 (2H, m), 2.46 (1H, m), 2.23 (1H, m), 1.73-1.79 (3H, m), 1.57-1.60 (1H, m), 1.35 and 1.41 (9H, s), 0.93 (3H, d, *J* = 6.5 Hz), 0.90 (3H, d, *J* = 6.5 Hz). ^13^C-NMR (125 MHz, DMSO-*d*_6_): δ 177.0, 171.6, 161.4, 155.4, 154.1, 138.3, 130.3, 124.9, 118.5, 105.9, 101.7, 99.7, 80.4, 55.5, 50.3, 40.6, 39.9, 28.3, 26.7, 24.8, 23.5, 21.8. HRMS (ESI): m/z [M+Na]^+^ calcd for C_26_H_37_N_5_NaO_6_ 538.2636; found 538.2642.

### *N*-((*S*)-1-(2-((*R*)-2-Chloro-2-fluoroacetyl)-2-(((*R*)-2-oxopyrrolidin-3-yl)methyl)hydrazineyl)-4-methyl-1-oxopentan-2-yl)-4-methoxy-1*H*-indole-2-carboxamide (10b)

A solution of **13b** (197 mg, 0.383 mmol), HCl/dioxane (4 M, 4.0 mL) was stirred for 50 min atroom temperature under argon. The reaction mixture was concentrated in vacuo, and the residue was dissolved to DMF (2 mL), then (*R*)-CFA (45.7% in 1,4-dioxane, 78.9 µL, 0.450 mmol), TEA (214 µL, 1.53 mmol), and COMU (253 mg, 0.591 mmol) was added at 0 °C. After 10 min, the reaction mixture was poured into saturated aqueous NaHCO_3_ and extracted with ethyl acetate. The organic layer was separated, washed with saturated brine, dried over MgSO_4_, filtrated, and concentrated in vacuo. The residue was purified by medium pressure column chromatography (Biotage® Sfär Silica HC D 20 µm 5 g, chloroform:methanol = 100:0 to 95:5) to give **10b** (29.7 mg, 15%) as a light brown solid.

^1^H-NMR (500 MHz, DMSO-*d*_6_): δ 11.60 (1H, s), 10.61 and 11.02 (1H, s), 7.74 and 7.76 (1H, br), 7.39 and 7.41 (1H, br), 7.11 (1H, dd, *J* = 8.0, 7.5 Hz), 7.03 (1H, d, *J* = 8.0 Hz), 6.26 and 6.88 (1H, d, *J* = 50.0 Hz), 6.51 (1H, d, *J* = 7.5 Hz), 4.32 and 4.48 (1H, br), 3.85 (1H, m), 3.40 (1H, m), 3.10-3.21 (2H, m), 2.24 and 2.60 (1H, m), 2.24 (1H, m), 1.77-1.82 (3H, m), 1.52 and 1.62 (1H, m), 0.97 (3H, d, *J* = 6.0 Hz), 0.93 (3H, d, *J* = 6.0 Hz). ^13^C-NMR (125 MHz, DMSO-*d*_6_): δ 176.3, 173.6, 166.2 (d, *J*_(C–F)_ = 25.0 Hz), 154.1,138.4, 129.7, 125.1, 118.5, 106.0, 102.2, 99.7, 90.8 (d, *J*_(C–F)_ = 241.3 Hz), 55.5, 51.3. 49.9, 39.1, 26.7, 24.9, 23.6, 21.6. HRMS (ESI): *m/z* [M+Na]^+^ calcdfor C_23_H_29_ClFN_5_NaO_5_ 532.1733; found 532.1704.

### *N*-((*S*)-1-(2-((*S*)-2-Chloro-2-fluoroacetyl)-2-(((*R*)-2-oxopyrrolidin-3-yl)methyl)hydrazineyl)-4-methyl-1-oxopentan-2-yl)-4-fluoro-1*H*-indole-2-carboxamide (11)

^1^H-NMR (500 MHz, DMSO-*d*_6_): δ 11.96 (1H, m), 10.75 (1H, s), 8.75 (1H, m), 7.72 and 7.80 (1H, br), 7.41and 7.43 (1H, br), 7.28 (1H, d, *J* = 8.0 Hz), 7.17 (1H, ddd, *J* = 13.0, 8.0, 5.0 Hz), 6.69 and 7.03 (1H, d, *J* = 50.0 Hz), 6.82 (1H, dd, *J* = 11.0, 8.0 Hz), 4.47 (1H, m), 3.86 and 4.01 (1H, m), 3.06-3.57 (3H, m), 2.48 and 2.63 (1H, m), 2.20 (1H, m), 1.61-1.83 (4H, m), 0.97 (3H, d, *J* = 6.0 Hz), 0.92 (3H, d, *J* = 6.0 Hz). ^13^C-NMR (125 MHz, DMSO-*d*_6_): δ 176.4 and 176.7, 171.5 and 173.0, 165.0 and 165.3 (d, *J*_(C–F)_ = 25.6 Hz), 161.8 and 162.8, 156.6 (d, *J*_(C–F)_ = 245.0 Hz), 139.5 (d, *J*_(C–F)_ = 10.0 Hz), 131.7, 124.6 (d, *J*_(C–F)_ = 7.5 Hz), 116.6 (d, *J*_(C–F)_ = 22.5 Hz), 109.4, 104.5 (d, *J*_(C–F)_ = 18.8 Hz), 99.8 and 99.9, 90.9 and 91.4 (d, *J*_(C–F)_ = 241.9 Hz), 66.4, 50.9 and 51.8, 49.0 and 49.4, 38.4, 26.4 and 26.6, 24.8, 23.4, 21.8. HRMS (ESI): *m/z* [M+Na]^+^ calcd for C_22_H_26_ClF_2_N_5_NaO_4_ 520.1534; found 520.1563.

### *N*-((*S*)-1-(2-Acetyl-2-(((*S*)-2-oxopyrrolidin-3-yl)methyl)hydrazineyl)-4-methyl-1-oxopentan-2-yl)-4-fluoro-1*H*-indole-2-carboxamide (12)

^1^H-NMR (500 MHz, DMSO-*d*_6_): δ 11.94 (1H, br), 10.51 and 10.72 (1H, br), 8.73 (1H, br-d), 7.68 (1H, br), 7.42 (1H, d, *J* = 1.0 Hz), 7.26 (1H, d, *J* = 8.0 Hz), 7.16 (1H, ddd, *J* = 13.0, 8.0, 5.0 Hz), 6.81 (1H, d, *J* = 10.5, 8.0 Hz), 6.81 (1H, dd, *J* = 10.5, 8.0 Hz), 4.48 (1H, br), 3.72-3.89 (1H, m), 3.10-3.50 (3H, m), 2.45-2.46 (1H, m), 2.17 (1H, br), 1.91 (3H, s), 1.70-1.83 (3H, m), 1.54-1.60 (1H, m), 0.96 (3H, d, *J* = 6.5 Hz), 0.91 (3H, d, *J* = 6.0 Hz). ^13^C-NMR (125 MHz, DMSO-*d*_6_): δ 176.6, 171.7 and 172.2, 171.3, 160.9, 156.2 (d, *J*_(C–F)_ = 244.3 Hz), 139.0 (d, *J*_(C–F)_ = 10.8 Hz), 131.4, 124.0 (d, *J*_(C–F)_ = 7.6 Hz), 116.2 (d, *J*_(C–F)_ = 21.8 Hz), 108.9 (d, *J*_(C–F)_ = 3.4 Hz), 104.0 (d, *J*_(C–F)_ = 18.1 Hz), 99.2, 56.0, 50.3. 48.0, 47.4, 26.7, 25.9, 24.4, 22.9, 21.4, 20.4. HRMS (ESI): *m/z* [M+Na]^+^ calcd for C_22_H_28_FN_5_NaO_4_ 468.2018; found 468.2061.

## Notes

### Competing Interest Statement

The authors have declared no competing interest.

